# Arsenic hexoxide has differential effects on cell proliferation and genome-wide gene expression in human primary mammary epithelial and MCF7 cells

**DOI:** 10.1101/2021.01.12.426459

**Authors:** Donguk Kim, Na Yeon Park, Keunsoo Kang, Stuart K. Calderwood, Dong-Hyung Cho, Ill Ju Bae, Heeyoun Bunch

## Abstract

Arsenic is reportedly a biphasic inorganic compound for its toxicity and anticancer effects in humans [1, 2]. Recent studies have shown that certain arsenic compounds including arsenic hexoxide (AS_4_O_6_; hereafter, AS6) induce programmed cell death and cell cycle arrest in human cancer cells and murine cancer models [3, 4]. However, the mechanisms by which AS6 suppresses cancer cells are incompletely understood. In this study, we report the mechanisms of AS6 through transcriptome analyses. In particular, the cytotoxicity and global gene expression regulation by AS6 were compared in human normal and cancer breast epithelial cells. Using RNA-sequencing and bioinformatics analyses, differentially expressed genes in significantly affected biological pathways in these cell types were validated by real-time quantitative polymerase chain reaction and immunoblotting assays. Our data show markedly differential effects of AS6 on cytotoxicity and gene expression in human mammary epithelial normal cells (HUMEC) and Michigan Cancer Foundation 7 (MCF7), a human mammary epithelial cancer cell line. AS6 selectively arrests cell growth and induces cell death in MCF7 cells without affecting the growth of HUMEC in a dose-dependent manner. AS6 alters the transcription of a large number of genes in MCF7 cells, but much fewer genes in HUMEC. Importantly, we found that the cell proliferation, cell cycle, and DNA repair pathways are significantly suppressed whereas cellular stress response and apoptotic pathways increase in AS6-treated MCF7 cells. Together, we provide the first evidence of differential effects of AS6 on normal and cancerous breast epithelial cells, suggesting that AS6 at moderate concentrations induces cell cycle arrest and apoptosis through modulating genome-wide gene expression, leading to compromised DNA repair and increased genome instability selectively in human breast cancer cells.

## INTRODUCTION

Cancer is one of the leading causes of human mortality; according to the World Health Organization (WHO), it ranked sixth among the top 10 causes of global death in 2016 (https://www.who.int/news-room/fact-sheets/detail/the-top-10-causes-of-death). Although a measurable number of cancers are both preventable and treatable, no anticancer drug or method has yet been developed to prevent and treat cancers effectively without affecting normal cells [5]. Over many years, researchers worldwide have made extensive and rigorous efforts to identify and develop such drugs. Breast cancer is the most common cancer in women. It is estimated that more than 508,000 women worldwide died from breast cancer in 2011 (WHO, https://www.who.int/cancer/detection/breastcancer/en/index1.html), and about 42,170 women have died from the disease so far in 2020 in the USA (American Cancer Society,https://www.cancer.org/cancer/breast-cancer/about/how-common-is-breast-cancer.html).The disease affects both developed and developing countries similarly[6]. The survival rates among breast cancer patients with the current remedies are low and vary by country, ranging from 40% to 80% [6]. The worldwide incidence has been increasing by 0.3% per year [7].

Current chymotherapy treatments for breast cancer can be categorized as selective estrogen receptor (ER) modulators, estrogen production inhibitors, and cell growth receptor modulators [8-10]. Most estrogen receptor modulators resemble the hormone estrogen and, although ineffective at triggering ER mediated transcription, bind to the estrogen receptor instead of the hormone to prevent breast cancer cells from proliferating[8]. Aromatase inhibitors block the production of estrogen [11].

Biological response modulators target specific proteins on the cell surface of breast cancer cells (e.g.HER2) to suppress their growth [12]. Yet because these drugs affect both normal and cancerous breast cells, side effects, including neuropathy, osteoporosis, infertility, lymphedema, and more, are inevitable.

Arsenic, a naturally forming element that often occurs in combination with sulfur and other metals, is an environmental contaminant found in drinking water [13]. The WHO and the International Agency for Research on Cancer categorize inorganic arsenic as carcinogenic, and provisional guidelines recommend a concentration below 10 μg/L [14]. However, it has been reported that about 50 countries around the world have drinking water with concentrations of inorganic arsenic over the recommended level[15, 16]. Exposure to inorganic arsenic over a prolonged period of time is reportedly linked to the occurrence of skin, bladder, and lung cancer [17].

Two inorganic arsenic compounds—arsenic trioxide and arsenic hexoxide (AS_4_O_6_; hereafter, AS6)— have already been used or developed as anticancer medications. Arsenic trioxide (AS_2_O_3_) has been developed into a commercially available cancer drug to treat acute promyelocytic leukemia (APL) with few and relatively mild adverse effects [1, 18]. It works by targeting diverse cellular pathways that lead to apoptosis and myeloid differentiation in APL, although its mechanisms have not been completely known [18]. Some studies suggest that arsenic trioxide treatment leads to apoptosis as a result of Jun N-terminal kinase suppression and collapse of mitochondrial transmembrane potentials to activate caspase 3 [19, 20]. Another study has proposed that arsenic trioxide at low concentrations between 0.1–0.5 μM promotes cell differentiation by degrading promyelocytic leukemia protein-retinoic acid receptor alpha (PML-RAR*α*) oncoprotein [21, 22] while inducing apoptosis at higher concentrations between1 and 2 μM [23]. It is interesting that arsenic trioxide and all-trans retinoic acid exhibit synergic effects to cure APL [24]. For example, in one study, 2-year event-free survival rates were 97% for an arsenic trioxide-all-trans retinoic acid group and 86% for an all-trans retinoic acid/chemotherapy group [25]. In addition, arsenic trioxide eradicates latent human immunodeficiency virus-1 (HIV-1) by reactivating latent provirus in CD4+ T cells and increasing immune responses in HIV-1 patients [26]. This study showed that arsenic trioxide treatment downregulates CD4 receptors and CCR5 co-receptors of CD4+ T cells that can interfere with viral infection and rebound [26].

AS6 is another arsenic compound that has been investigated and developed as an anticancer drug [4, 27-29]. It has been suggested that AS6 has distinctive anticancer effects from arsenic trioxide [30]. Two recent studies indicated that AS6 induces apoptosis, G2/M cell cycle arrest, and autophagy in colon cancers [4]. Using targeted approaches, the studies suggested that AS6 induces cell death through effects on the p38 MAPK pathways in a colon cancer cell line, SW620 cells [4]. Another study reported that AS6 inhibits NF-*κ*B to stimulate TNF-*α*-induced cell death in MCF7 cells [31]. In spite of these efforts, the effects and anticancer mechanisms of AS6 on human cells are not completely understood and require unbiased approaches to identify them. In addition, it is important to determine whether AS6 differentially affects normal and cancerous cells and to what extent.

To address these questions, we have compared the cytotoxicity of AS6 in normal and cancerous breast epithelial cells, human mammary epithelial normal cells (HUMEC), and Michigan Cancer Foundation 7 (MCF7) cells. An interesting finding was that MCF7 cells were much more susceptible to AS6 than HUMEC, and some major cell cycle factors were differentially regulated in these cells by AS6. Therefore, we further investigated the impact of AS6 on genomic expression and used RNA-sequencing (RNA-seq) analysis to analyze the changes in expression of all protein-coding and non-coding genes in these cells with or without AS6 treatment. Strikingly, AS6 altered the transcription of a large number of genes in MCF7 cells whereas much fewer genes were differentially expressed in HUMEC in the same condition. Our bioinformatics analyses have identified a number of cellular pathways that were significantly impacted by AS6 in MCF7 cells. The downregulated pathways included cell cycle progression, DNA replication, and DNA repair whereas the upregulated pathways involved apoptosis and stress-response. Furthermore, we validated these genomic data through the real-time PCR and immunoblotting analyses to quantify the expression of several critical genes at the RNA and protein levels. Our intriguing results suggest that AS6 treatment at concentrations between 0.1 and 1 μM increases cellular stress and genomic instability, ultimately inducing cell cycle arrest and apoptosis in MCF7 cells, but not in HUMEC. Our results therefore, provide the essential and fundamental understanding of the cytotoxicity, anticancer effects, and genome regulation of AS6 for the future evaluations and applications in the environmental and medical fields.

## MATERIALS & METHODS

### Chemicals

Arsenic hexoxide (AS_4_O_6_, AS6) used in this study was provided by Chemas Co., LTD (Seoul, South Korea). AS6 was invented by Ill Ju Bae and Zenglin Lian and was manufactured by the company. The chemical properties and purity of AS6 were validated through the analytical chemical methods. AS6 has been patented for treating breast cancer under a United States patent number US 10,525,079 B2 since January 7, 2020. AS6 was provided as a 2.5 mM stock solution dissolved in water.

### Cell Culture

Primary mammary normal epithelial cells (HUMEC, ATCC PCS-600-010) and MCF7 (ATCC HTB-22) were purchased from the American Type Culture Collection (ATCC, USA). The cells were cultured at 37°C in a 5% CO_2_ incubator. HUMEC were maintained in HuMEC Ready Medium containing HuMEC Basal Serum Free Medium, HuMEC Supplement, and bovine pituitary extract (Cat # 12752-010, Thermo Fisher Scientific, USA). MCF7 cells were cultured in DMEM (Cat # 10013CV, Corining, USA) containing 10% fetal bovine serum (FBS, Gibco, USA) and 1% penicillin/streptomycin (P/S, Thermo Fisher, USA).

### Cytotoxicity test

HUMEC and MCF7 cells were grown to 70–80% confluence in a 10 cm dish before splitting into a 96 well plate. Approximately 4 × 10^3^ cells were seeded in each well and AS6 was applied according to indicated concentrations. After 24–72 h incubation, 10% (v/v) of water soluble tetrazolium salt (WST, DoGen Inc., South Korea) was added to each well, following the manufacturer’s instruction. Intensity of orange color developed from the enzyme-substrate reaction was measured at 450 nm using spectrophotometry (BMG Labtech, Germany). Cell images were taken by Olympus IX-71 microscope (Olympus, Japan) equipped with objective lenses (Olympus LUCPLFLN20X, Olympus, Japan), a camera (Olympus XM10, Olympus, Japan), and a light source (Olympus TH4-200, Olympus, Japan). Images were acquired, using CellSens Standard Imaging software (Olympus, Japan).

### RNA-seq

Cell culture and RNA preparation were performed as described in [32]. HUMEC and MCF7 cells were grown to 60-70% confluence in 6-well plates and the media were exchanged with the fresh complete media including AS6 at final concentration of 0.5 µM or H_2_O only as an untreated control. After 50 h incubation, the cells were washed with cold PBS once and scraped. The cells were washed again with cold PBS twice before extracting total RNA molecules using Qiagen RNeasy Kit (Qiagen, Germany). The cDNA construction and RNA sequencing were commercially performed by Omega Bioservices (https://omegabioservices.com,USA) using the manufacturer’s typical procedure as previously reported (https://www.ncbi.nlm.nih.gov/geo/query/acc.cgi?acc=GSM471292)[33]. The RNA concentration and integrity were assessed using Nanodrop 2000c (Thermo Fisher Scientific, USA) and Agilent 2200 Tapestation instrument (Agilent Technologies, USA). One microgram of total RNA was used to prepare Ribo-Zero RNA-Sequencing (RNA-Seq) libraries. Briefly, ribosomal RNA (rRNA) is removed using biotinylated, target-specific oligos combined with Ribo-Zero rRNA removal kit (Illumina, USA). After purification, the RNA is fragmented into small pieces using divalent cations under elevated temperature. First-strand cDNA syntheses were performed at 25°C for 10 minutes, 42°C for 15 minutes and 70°C for 15 minutes, using random hexamers and ProtoScript II Reverse Transcriptase (New England BioLabs, USA). In a second strand cDNA synthesis the RNA templates were removed, and a second replacement strand was generated by incorporation dUTP (in place of dTTP, to keep strand information) to generate ds cDNA. The blunt-ended cDNA was cleaned up from the second strand reaction mix with beads. The 3’ends of the cDNA were then adenylated and followed by the ligation of indexing adaptors. PCR (15 cycles of 98°C for 10 seconds, 60°C for 30 seconds and 72°C for 30 seconds) were used to selectively enrich those DNA fragments that have adapter molecules on both ends and to amplify the amount of DNA in the library. The libraries were quantified and qualified using the Agilent D1000 ScreenTape on a 2200 TapeStation instrument. The libraries were normalized, pooled and subjected to cluster and pair-read sequencing was performed for 150 cycles on a HiSeqX10 instrument (Illumina, USA), according to the manufacturer’s instructions.

### Bioinformatics

Sequenced reads were trimmed to remove portions of poor sequenced quality (Phred score < 20) and/or contaminated adapter sequences using Trim Galore (version 0.6.4; https://www.bioinformatics.babraham.ac.uk/projects/trim_galore/). Trimmed reads were aligned to the human reference genome (hg38 assembly) using STAR (version 2.7.3a) [34] with default parameters. The abundance of transcripts was quantified using StringTie (version 2.0.6) [35] by means of transcripts per million (TPM). Then, differentially expressed genes (DEGs) were identified using DESeq2 (version 1.24.0) [35] with an adjusted p-value cutoff of 0.05. Among the identified DEGs, transcripts showing less than two fold-change between comparisons and average TPM value of 1 across samples were further discarded. Heatmaps were generated using the Morpheus web application (https://software.broadinstitute.org/morpheus/) with the min-max normalization.

Hierarchical clustering of genes in the heatmaps was conducted by the average linkage algorithm with the one-minus pearson correlation metric. GO and PPI analyses of DEGs were performed using Metascape [36] (https://metascape.org).

### Real-time PCR

Cell culture and RNA preparation for real-time PCR were performed as described in [37]. HUMEC and MCF7 cells were grown to 60–70% confluence in six well plates and were replaced with the fresh media before applying AS6 to the target concentrations. After 48–72 h incubation, the cells were washed with cold PBS once and scraped. The cells were washed again with cold PBS twice before the total RNA molecules were extracted using the Qiagen RNeasy kit. cDNAs were constructed from 136–600 ng of the collected RNAs using ReverTra Ace qPCR RT Master Mix (Toyobo, Japan). cDNA was analyzed through qPCR using SYBR Green Realtime PCR Master Mix (Toyobo, Japan), according to the manufacturer’s instructions (QuantStudio3 Real-Time PCR System, Applied Biosystems, Thermo Fisher Scientific, USA). The thermal cycle used was 95°C for 1 min as pre-denaturation, followed by 45 cycles of 95 °C for 15 s, 55 °C for 15 s, and 72 °C for 45 s. The primers used for the experiments were purchased from Integrated DNA technology (USA) and are summarized in **Table S1**.

### Western blot

MCF7 cells were grown in 6 well plates for Western blots and were washed with cold PBS twice and scraped in RIPA buffer (Cell signaling, USA). Protein concentration in each sample was measured through Bradford assay using Bio-Rad Protein Assay Dye Reagent Concentrate (Bio-Rad, USA) and spectrophotometry at 595 nm (BMG Labtech, Germany). From the measured protein concentration, a total of 20 µg of proteins per sample was loaded on 6–12% SDS-polyacrylamide gels, and transferred to PVDF or nitrocellulose membrane (Bio-Rad, Thermo Fisher Scientific, USA). After blocking with 4– 5% skim milk (MBcell, South Korea) in TBST including 25 mM Tris (Sigma, USA), 140 mM NaCl (Thermo Fisher Scientific, USA), and 0.05% Tween 20 (Sigma, USA), the membrane was probed with indicated primary antibodies. Anti-BUB1B (A300-386A) and anti-BRCA1 (A300-000A) antibodies were purchased from Bethyl laboratories (USA); anti-PLK1 (sc-17783), anti-CDC25A (sc-7389), anti-CDC20 (sc-13162), anti-CHEK1 (sc-8408), anti-CDKN1A (sc-6246), anti-HSP90 (sc-69703) and anti-HSP70 (sc-32239) antibodies were purchased from Santa Cruz Biotechnology (USA); anti-CCNB1 (#4135) and anti-HIF1*α* (#14179) antibodies were purchased from Cell Signaling Technology (USA); anti-ATM (ab78) and anti-MCM4 (ab4459) antibodies were purchased from Abcam; anti-ACTA1 (ACTIN, MAB1501) was purchased from Sigma Aldrich (USA). Antibodies were validated for Western blotting by the manufacturers. Antibody were diluted to 1:300–1:2000 in blocking solution for usage as recommended by the manufacturers. For protein detection, the membranes were incubated with HRP-conjugated mouse and rabbit secondary antibodies (Cell Signaling Technology, 7076S and 7074S) and signals were detected with EzWestLumi Plus or Western blotting Luminol Reagent (ATTO, WSE-7120L, Japan; sc-2048, Santa Cruz Biotechnology, USA). Image J software (National Institute of Health, USA) was used to measure the band intensity of immunoblotting.

### Statistical analyses

Statistical analyses were performed as described in [37]. One-way ANOVA, followed by a Tukey’s HSD test, were used to determine differences in toxicity percentages (*P* < 0.05) (SAS for Window release 6, SAS Institute). Log-probit regression was used to determine the LC_50_ based on corrected mortality from different chemical concentrations (SAS for Windows release 6, SAS Institute). All analyses were performed in SAS version 9.4. One- or two-way ANOVA was used to determine differences (*P* < 0.05) for mRNA quantification and RNA-seq. Graphs were drawn using Prism 8 (GraphPad, Inc., San Diego, CA, USA).

## RESULTS

Although AS6 reportedly induces cell death in colon and breast cancer cells [4, 31], preliminary clinical data suggested that it might have few and mild side effects in humans (CA Patent #: CA2298093C). These findings suggest that the compound affects normal and cancer cells differentially, inducing potentially more serious cytotoxicity in the cancer cells. Therefore, we first tested whether the cytotoxicity of AS6 differs between the normal and cancerous mammary epithelial cells, HUMEC and MCF7 cells. Cells were treated with AS6 at concentrations of 0, 0.5, 10, 50, 100, and 200 μM and were monitored for viability at 24h and 48 h with the WST assay (**Fig. 1A**). The results clearly showed that AS6 was toxic to both cell types within 24 h at concentrations over 10 μM (**Fig. 1B**; **Supplementary Fig. 1A**). Both types of cells, when treated with 10 μM AS6, had survival rates less than 10% at 48 h (**Fig. 1B**; **Supplementary Fig. 1B**), which indicates the significant cytotoxicity of AS6 to human mammary cells at this concentration. Interestingly however, the treatment with 0.5 μM AS6 resulted in differential responses between HUMEC and MCF7 cells: HUMEC increased in population, whereas MCF7 cell numbers decreased at 48 h (**Fig. 1B**; **Supplementary Fig. 1B**). Microscopic observation suggested that HUMEC grew normally, whereas the growth of MCF7 cells was markedly compromised with 0.5 μM AS6 at 48 h (**Supplementary Fig. 1B**).

**Figure 1.**
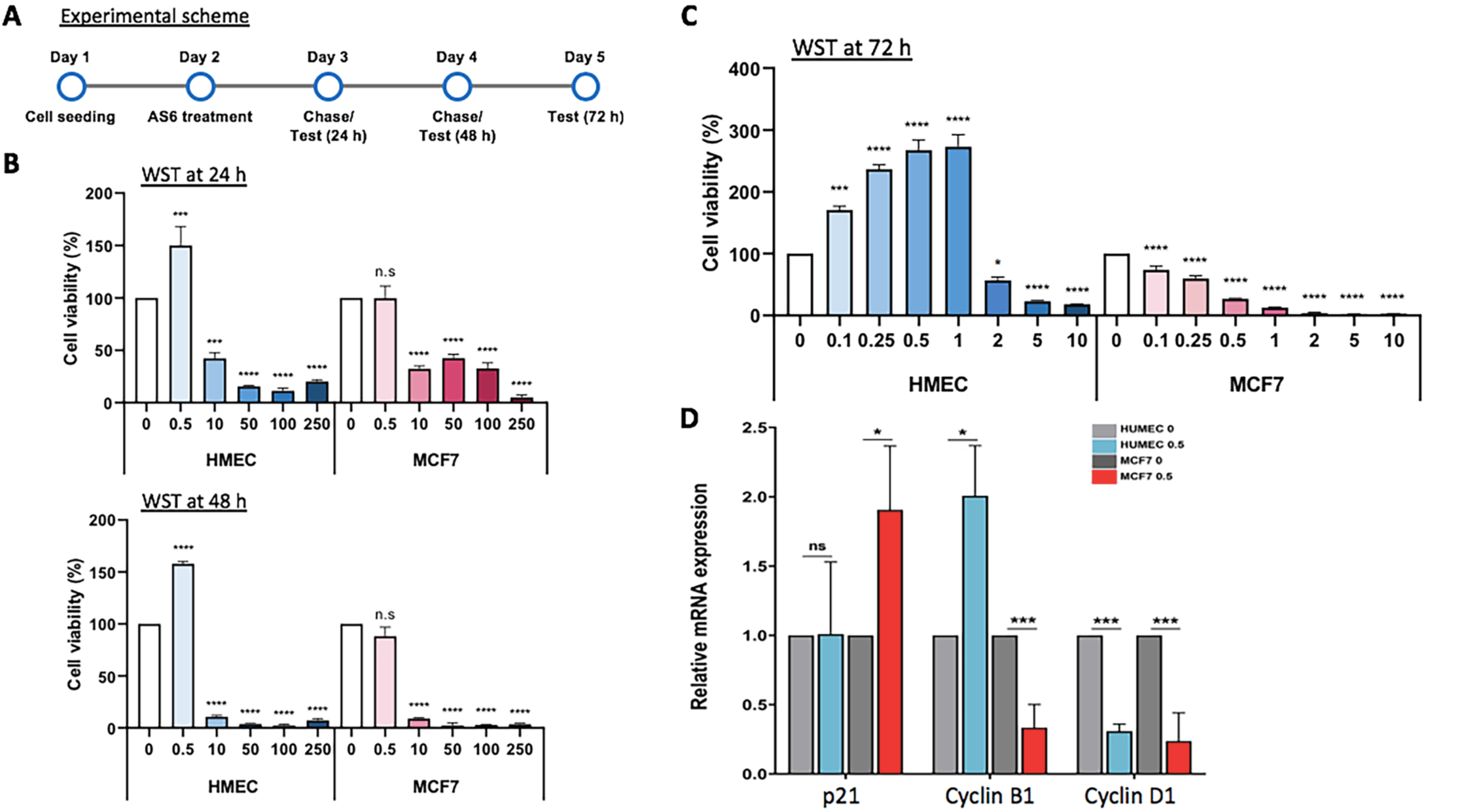
AS6 has markedly differential cytotoxic effects in HUMEC and MCF7 cells. **(A)** Timeline for cytotoxicity assays. A brief experimental scheme from cell seeding to WST assays. **(B)** WST data with 0–250 μM AS6 (X-axis) treated for 24 h (top) and 48 h (bottom) in HUMEC and MCF7 cells. *n* = 8; throughout the legends, n indicates the number of biological replicates. Error bars show standard deviations (s.d.). n.s, not significant; * * * P < 0.001; * * * * P < 0.0001. **(C)** WST data with 0–10 μM AS6 (X-axis) treated for 72 h in HUMEC and MCF7 cells. *n* = 8; error bars show s.d.; * P < 0.05; * * * P < 0.001; * * * * P < 0.0001. **(D)** RT-qPCR results of the quantification of *p21, cyclin B1*, and *cyclin D1* mRNAs in AS6-treated HUMEC and MCF7 cells. *ACTIN* was used as a reference gene for normalization. *n* = 3; error bars show s.d.; * P < 0.05; * * * P < 0.005.

Next, we investigated AS6 cytotoxicity to HUMEC and MCF7 cells at lower concentrations surrounding 0.5 μM: 0, 0.1, 0.25, 0.5, 1, 2, 5, and 10 μM. Cells were treated with AS6 for 72 h, and then the viability of cells was measured with WST assay. The results showed a dose-dependent cell death in MCF7 cells (**Fig. 1C**; **Supplementary Fig. 1C**). By contrast to this, HUMEC showed a biphasic effect dependent on the concentration of AS6: AS6 up to 1 μM did not interfere with cell growth, and even seemed to stimulate it, whereas concentrations over 2 μM did interfere with growth (**Fig. 1C**; **Supplementary Fig. 1C**). For example, the treatment with 1 μM AS6 for 72 h increased the density of viable cells to 278% in HUMEC but decreased cell density to 27% in MCF7 cells, about a 73% reduction (**Fig. 1C**; **Supplementary Fig. 1C**). It should be noted that the changes in the cell growth and morphology mediated by AS6 were microscopically clear and dramatically distinct (**Supplementary Fig. 1C**), which suggests a different susceptibility to AS6, between the HUMEC and MCF7 cells. Indeed, the LC_50_ values derived from the experiments were 8.8 times higher value for MCF7 cells than HUMEC, at 0.26 μM versus 2.29 μM, respectively (**Table 1**). This demonstrates that the malignant MCF7 cells are more susceptible to AS6 than HUMEC.

**Table 1.** LC_50_ of AS6 for HUMEC and MCF7 cells

| Cell type | LC<sub>50</sub>(μM) | 95% CI (lower – upper) | Slope ± SEM | X<sup>2</sup> (df) |
| --- | --- | --- | --- | --- |
| MCF7 | 0.26 | 0.13 – 0.40 | 1.6228 ± (0.2450) | 20.29 (5) |
| HUMEK | 2.29 | n/a | 1.6263 ± (0.4754) | 2.97 (1) |

CI, confidence interval; the LC<sub>50</sub> value was calculated using percentage mortality; n/a, no confidence interval observed and therefor no probit analysis performed; SEM, standard error of mean

To understand the effect of AS6 on gene expression, we monitored the transcription of critical cell cycle regulatory genes *cyclin B1, cyclin D1*, and *p21*. HUMEC and MCF7 cells were incubated with AS6 for 48 h before the reverse transcription quantitative PCR analysis (RT-qPCR). In line with the differential cytotoxicity of AS6 to HUMEC and MCF7 cells, expression of *cyclin B1* increased in HUMEC and decreased in MCF7 cells (**Fig. 1D**). *Cyclin B1* decreased significantly to less than 40% of the untreated control, which suggests a possible disruption of the G_2_–M cell cycle transition [38] in MCF7 cells (**Fig. 1D**). By contrast, *cyclin D1* was reduced in both HUMEC and MCF7 cells, suggesting a delayed G_1_–S transition [39] upon the treatment with AS6 (**Fig. 1D**). Interestingly, the mRNA expression level of p21 was increased in MCF7 cells but was unchanged in HUMEC (**Fig. 1D**). p21 is a potent cyclin-dependent kinase inhibitor that arrests G_1_, S, and G_2_ progression by interfering various CDKs including CDK1, 2, 4, and 6 [40]. We conclude that AS6 inhibits cell cycle progression through the G_1_, S, and G_2_ phase more extensively in MCF7 cells than it does in HUMEC.

We were prompted by these findings to further understand the impact of AS6 on the genome-wide gene expression in HUMEC and MCF7 cells using the RNA-seq analysis. Briefly, cells were treated with AS6 at a final concentration of 0.5 μM for 50 h, and the total RNA was collected in triplicates. To enhance the mRNA and ncRNA coverage, we depleted rRNAs before sequencing the collected RNA pool. A total of 81702 and 91152 protein-coding and non-protein coding genes in HUMEC and MCF7 cells were successfully sequenced (**Supplementary Fig. 2A**; **Supplementary Data 1**). The number of the genes, whose expression was increased or decreased more than 2-fold with statistical significance (|log_2_fold-change| > 1, p value < 0.05) when compared to untreated control, was 1233 and 7374 in HUMEC and MCF7 cells, respectively (**Fig. 2A, B**; **Supplementary Data 1**). Of these, 1059 and 5297 genes were differentially expressed more than 4-fold in HUMEC and MCF7 cells, respectively (**Fig. 2B**; **Supplementary Data 2**). The heat maps of the differentially expressed genes (|log_2_fold-change| > 1, p value < 0.05, n = 1233 and 7374 in HUMEC and MCF7 cells, respectively) clearly showed that many more genes, exceeding six times, were significantly affected in MCF7 cells than HUMEC by AS6 (**Fig. 2A, B**; **Supplementary Data 3**). All and differentially expressed genes (|log_2_fold-change| > 1, p value < 0.05) in HUMEC and MCF7 cells were compared in box plots (**Fig. 2C**). The genes that were up- and down-regulated (|log_2_fold-change| > 2, p value < 0.05; n= 599 and 460, respectively) in HUMEC were categorized by Gene Ontology (GO) analysis. The upregulated genes were involved in membrane trafficking and assembly, cell cycle transition, stress response, and DNA double strand break repair by homologous recombination (n= 599; **Fig. 2D**). A protein-protein interaction (PPI) network was constructed to identify the correlation among these differentially expressed protein-coding genes (**Fig. 2E**; **Supplementary Fig. 3A**). The downregulated genes were constituents of Ras signal transduction, oxidative stress response, and apoptosis pathways (n= 460; **Fig. 2F**). A PPI analysis of the downregulated protein-coding genes showed the networks and the degree of correlations of these genes (**Fig. 2G; Supplementary Fig. 3B**).

**Figure 2.**
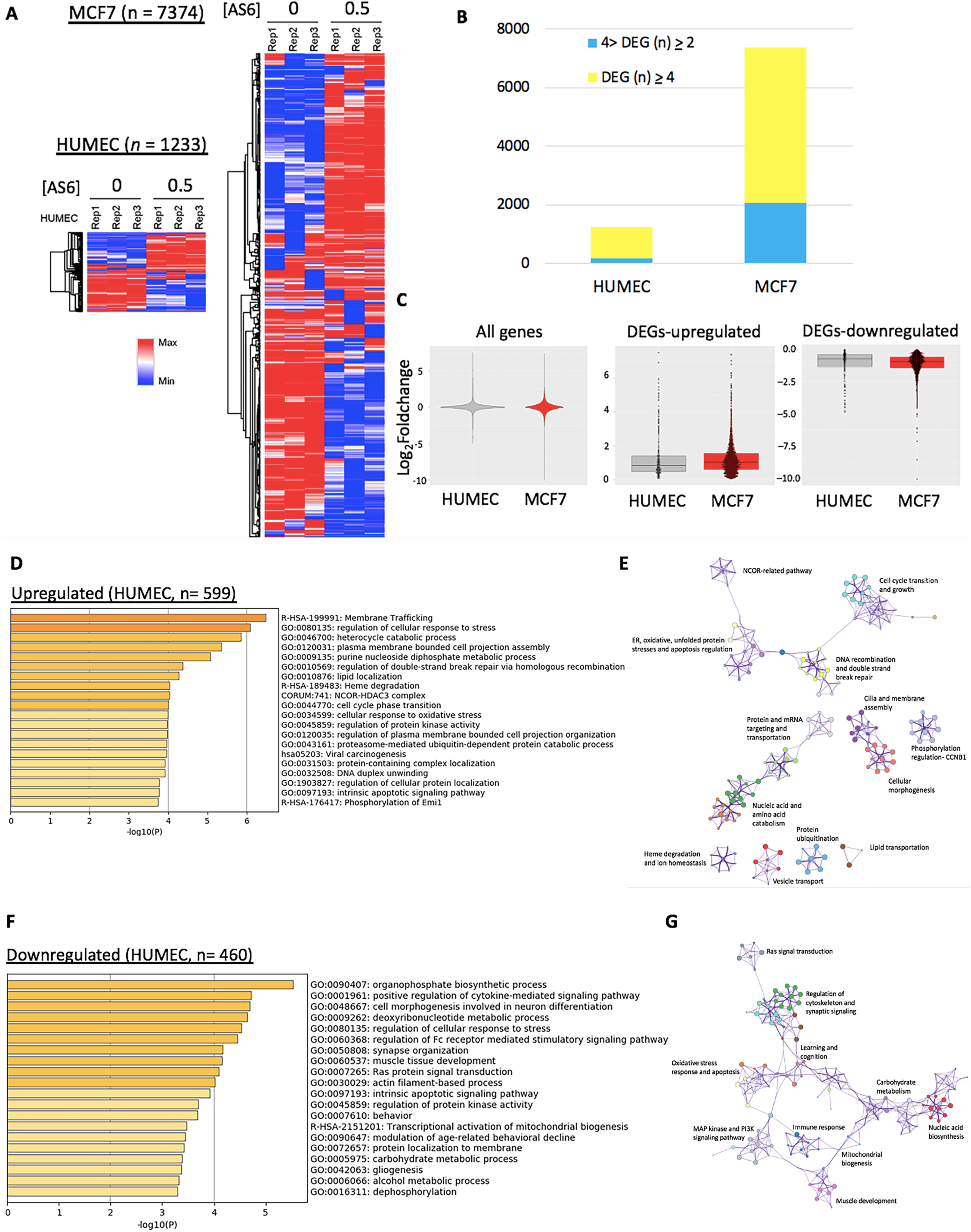
AS6 effects on genome-wide gene expression in HUMEC and MCF7 cells. **(A)** Heatmaps of differentially expressed genes upon AS6 treatment in HUMEC (left; *n* = 1233) and MCF7 cells (right; *n* = 7374). **(B)** Comparison of differentially expressed genes (foldchange > 4 in yellow) upon AS6 treatment between HUMEC and MCF7 cells displaying a significant difference (*n* = 1059 vs 5297, respectively) in responding to AS6 in these two cell types. **(C)** Box plots of all (left, *n* = 81707 and 91152 in HUMEC and MCF7 cells respectively) and differentially expressed genes [DEGs; up-(middle; *n* = 599 and 3772 in HUMEC and MCF7, respectively) and down-regulated (right, *n* = 634 and 3602 in HUMEC and MCF7, respectively)] comparing HUMEC and MCF7 cells. **(D)** GO analysis of upregulated genes in AS6-treated HUMEC. In all GO analysis tables, the X-axis is –log _10_P-value. **(E)** PPI analysis showing the relatedness of upregulated protein-coding genes in AS6-treated HUMEC. **(F)** GO analysis of downregulated genes in AS6-treated HUMEC. (**G**) PPI analysis showing the relatedness of downregulated protein-coding genes in AS6-treated HUMEC.

The genes in MCF7 cells, whose mRNA levels increased or decreased (|log_2_fold-change| > 1, p value < 0.05; n= 2815 and 2482) as a result of AS6 treatment were subjected to GO and PPI analyses (**Fig. 3A–D**). We observed some common pathways affected by AS6 between HUMEC and MCF7 cells including membrane trafficking and stress response (**Fig. 2D**; **Supplementary Fig. 3A**). On the other hand, unfolded protein response, exocytosis, apoptosis, hypoxia response, and ER stress response pathways were uniquely and significantly increased in MCF7 cells but not in HUMEC (**Fig. 2D–G, 3A–D**). **Figures 3B** and **Supplementary Fig. 3C** summarize the PPI network of those genes that were increased, compared to the untreated control. Strikingly, the genes downregulated upon AS6 treatment (n= 2482) greatly affected the cell cycle including mitotic nuclear division, cell division, cell cycle processes, microtubule-based processes, and G_2_/M phase transition (**Fig. 3C, D**). Other categories are closely related to cell growth and the cell cycle included DNA repair, telomere organization, the centrosome cycle, and regulation of chromosome organization (**Fig. 3C, D**). PPI analysis intriguingly showed the close relationship of downregulated protein-coding genes for the cell cycle progression/cell growth and genome integrity (**Fig. 3D**; **Supplementary Fig. 3D**). These results suggested that AS6 inhibits the cell cycle progression at the level of gene expression, resulting in the reduced cell viability as shown in the cytotoxicity analyses (**Fig. 1B, C**). **Tables 2–3** summarize and compare the pathways and representative protein-coding genes differentially affected by AS6 with statistical significance in HUMEC and MCF7 cells.

**Figure 3.**
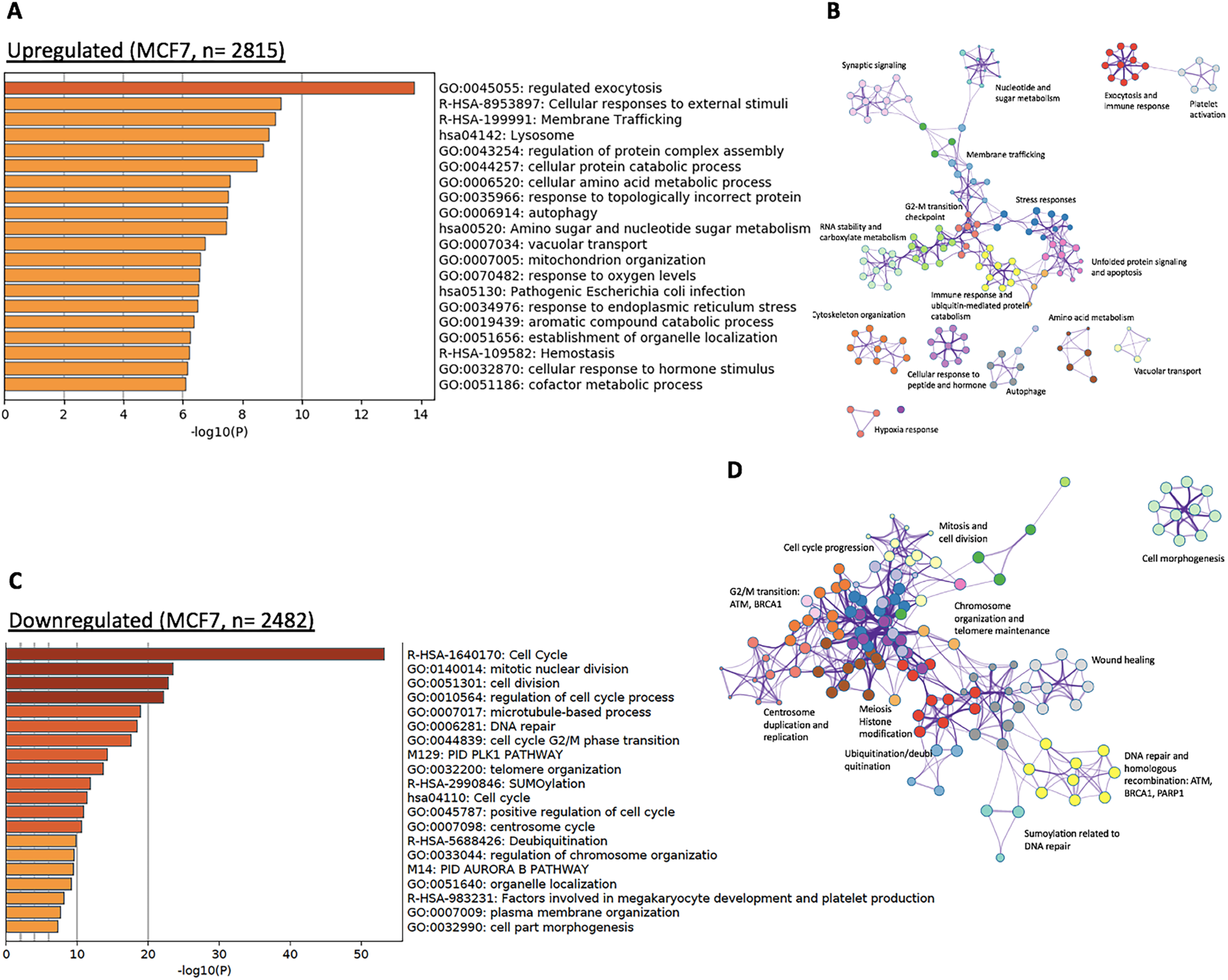
AS6 significantly disturbs critical biological pathways in MCF7 cells. **(A)** GO analysis of upregulated genes in AS6-treated MCF7 cells. **(B)** PPI analysis showing the relatedness of upregulated protein-coding genes in these cells. **(C)** GO analysis of downregulated genes in AS6-treated MCF7 cells showing a severe impairment of cell cycle and growth. **(D)** PPI analysis showing the marked relatedness of downregulated protein-coding genes in AS6-treated MCF7 cells.

**Table 2.** Biological pathways that are affected by AS6 in HUMEC and MCF7 cells

| **HUMEK** |  |
| --- | --- |
| *Upregulated pathways* | Membrane trafficking and assembly |
|  | Cellular responses to stresses |
|  | Nucleotide and protein catabolic pathways |
|  | Double strand DNA break repair through homologous recombination |
|  | Heme degradation |
|  | NCOR-HDAC3 complex |
|  | Cell cycle transition |
|  | Regulation of protein kinases |
| *Downregulated pathways* | Organophosphate biosynthetic process |
|  | Cytokine-mediated signaling pathway |
|  | Deoxyribonucleotide metabolic pathway |
|  | Muscle tissue development |
|  | Ras protein signaling transduction |
|  | Intrinsic apoptotic pathway |
|  | Transcriptional activation of mitochondrial biogenesis |
|  | Carbohydrate and alcohol metabolic pathways |
| **MCF7** |  |
| *Upregulated pathways* | Exocytosis |
|  | Cellular responses to external and internal stimuli and stresses |
|  | Membrane trafficking |
|  | Protein catabolic processes |
|  | Autophagy |
|  | Amino-sugar metabolism |
|  | Vacuolar transport |
|  | Mitochondria organization |
| *Downregulated pathways* | Cell cycle |
|  | Mitosis and G2/M transition |
|  | DNA repair |
|  | Telomere organization |
|  | Sumoylation |
|  | Centrosome cycle |
|  | Deubiquitination |
|  | Regulation of chromosome organization |
|  | Aurora B pathway |
|  | Membrane organization |
|  | Cellular morphogenesis |

**Table 3.**
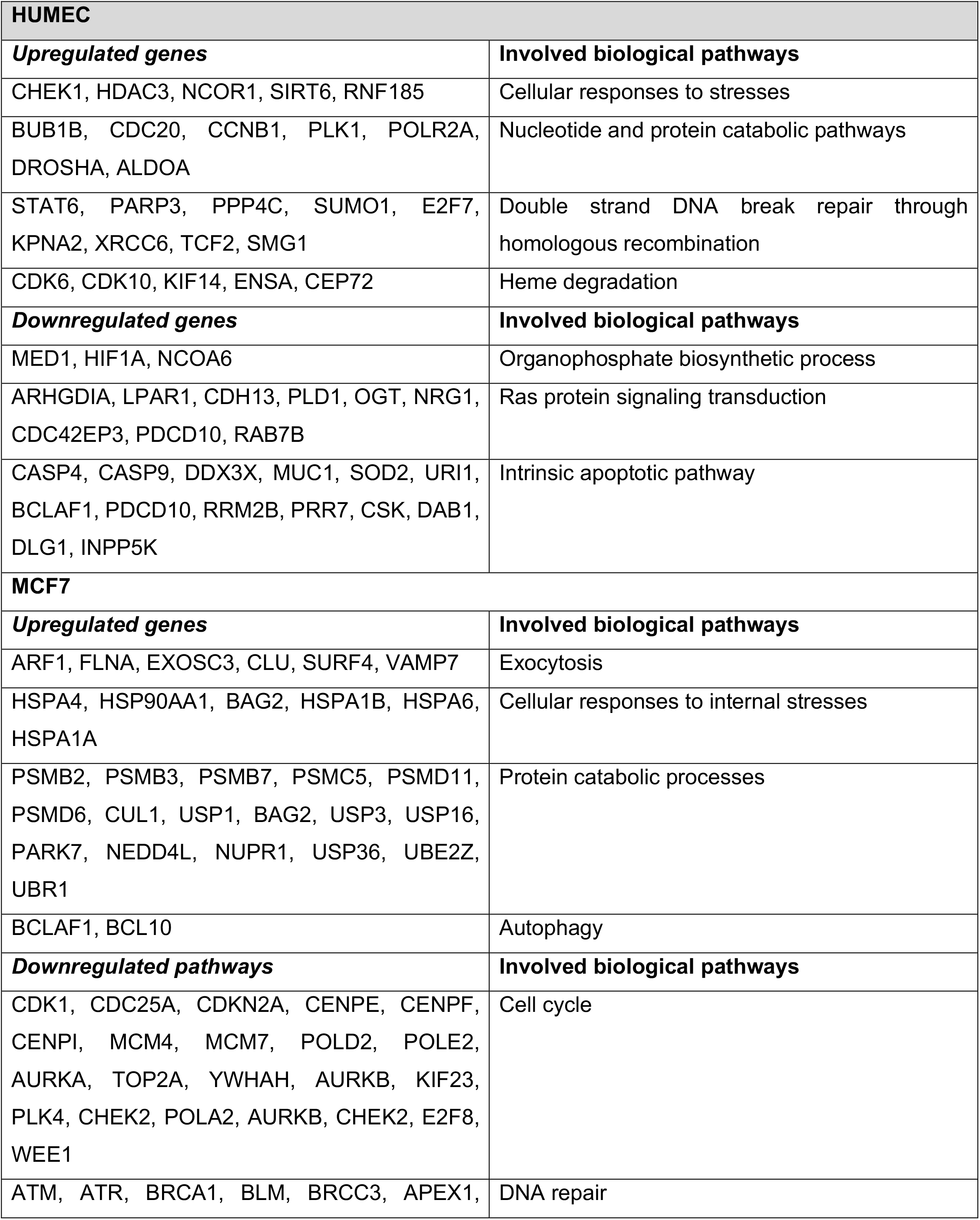
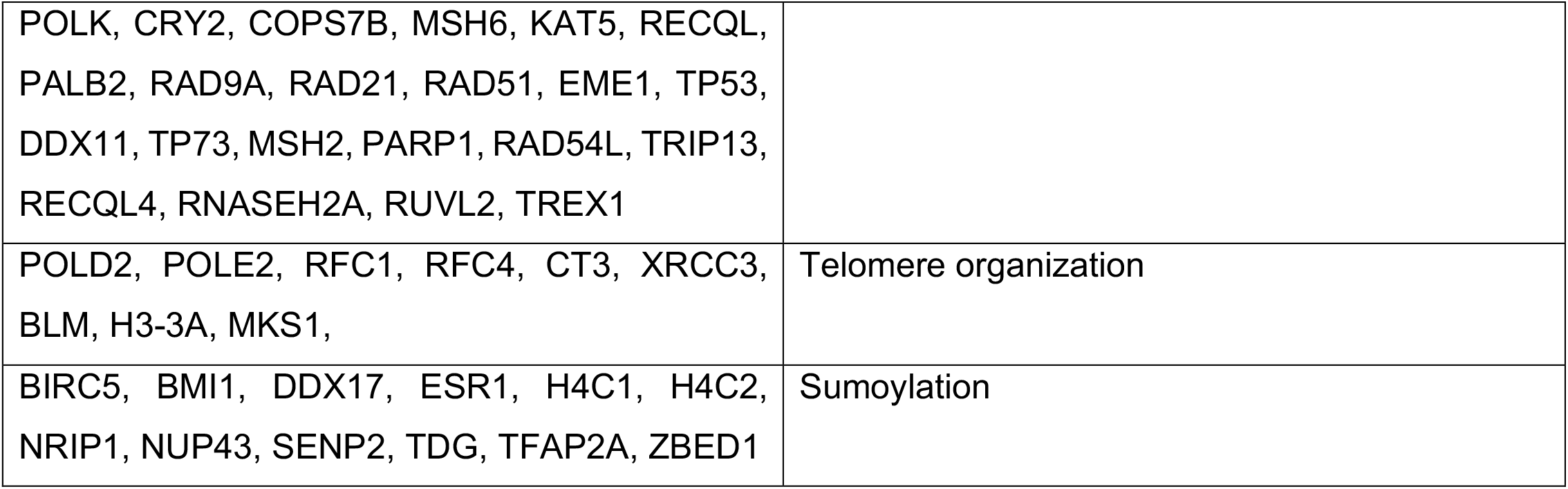
Representative protein-coding genes significantly affected by AS6 in HUMEC and MCF7 cells.

The RNA-seq and GO analyses suggested that modest concentrations of AS6 induced more deleterious cellular stresses in MCF7 cells than HUMEC (**Fig. 2D, 3A**). In addition, the analyses indicated that AS6 treatment resulted in the impairment of DNA repair function, the arrest of cell cycle progression, and the activation of apoptosis, uniquely in MCF7 cells, but not in HUMEC (**Fig. 2D–G, 3A–D**). We interpret this to mean that MCF7 cells and HUMEC have profoundly different sensitivity to AS6 at concentrations below 1 μM. It is plausible that MCF7 cells undergo apoptosis because increased genomic instability halts cell cycle progression. We conjecture that this genomic instability may be attributed to impaired DNA repair function in AS6-treated MCF7 cells.

The expression of a few critical protein-coding genes that were significantly affected by AS6 and are involved in cellular stress, DNA repair, cell cycle progression, and apoptosis was validated through real-time PCR and Western blotting. HUMEC and MCF7 cells were treated with AS6 at either 0.25 or 0.5 μM for 72 h and total RNAs or proteins were extracted. RNA-seq and bioinformatics analyses indicated that RNA expression of representative DNA repair proteins for DNA double strand break, including ataxia-telangiectasia mutated (*ATM*) and breast cancer susceptibility 1 (*BRCA1*) [41], was notably reduced in AS6-treated MCF7 cells (**Fig. 4A**; **Supplementary Fig. 4A**; **Table 3**). These proteins are essential for homology-directed DNA double strand break repair (HDR) [42, 43], which is error-free [43, 44]. Because of the enhanced genomic fidelity that can be achieved by HDR compared to non-homologous end joining, these repair proteins are considered to be critical, and their mutations or malfunctions have been implicated in various cancers [45, 46]. In particular, *ATM* and *BRCA1* mutations are strongly linked to breast cancers [47, 48]. An inherited mutated copy of ATM confers an increased risk of developing breast cancer as well as pancreatic, prostate, colon, and other cancers [49]. *BRCA1* is a potent tumor suppressor gene, whose mutations increase the risk for breast and ovarian cancers [46]. Such mutations also increase the risk for other cancers including pancreatic cancer, prostate cancer, and cervical cancers. In addition, BRCA1 deficiency can lead to an accumulation of DNA double strand breaks in genes activated by estrogen receptor [50]. p53 (TP53) is a critical genome guardian protein that senses the status of DNA breaks and genome integrity, leading to DNA repair, cell cycle arrest, or apoptosis [51]. Real-time PCR using AS6-treated HUMEC and MCF7 cells indicated reduced *BRCA1* and *ATM* expression in MCF7 cells but not in HUMEC (**Fig. 4B**). In particular, *BRCA1* mRNA levels decreased more than 80% in MCF7 cells compared to the control. *ATM* expression was downregulated in both HUMEC and MCF7 cells but more significantly and extensively in MCF7 cells (**Fig. 4B**). Immunoblotting results demonstrated that the protein levels of BRCA1 and ATM as well as p53 (for p53, also see below) were dramatically reduced by AS6 treatment in MCF7 cells (**Fig. 4C**; **Supplementary Fig. 4A**). These data suggest malfunctions in HDR and the sensing of DNA breaks and an increase in genome instability in MCF7 cells that are treated with AS6.

**Figure 4.**
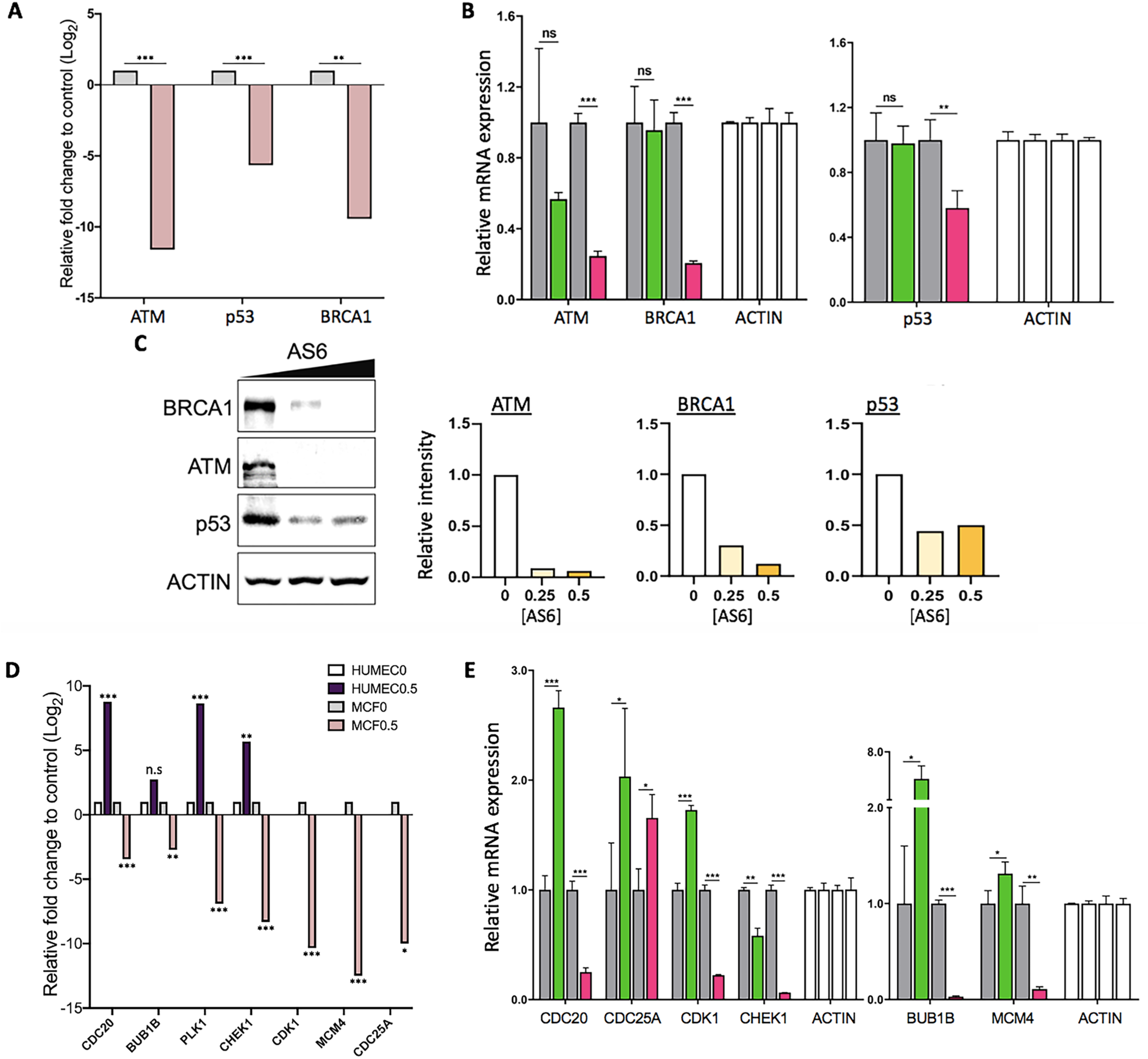
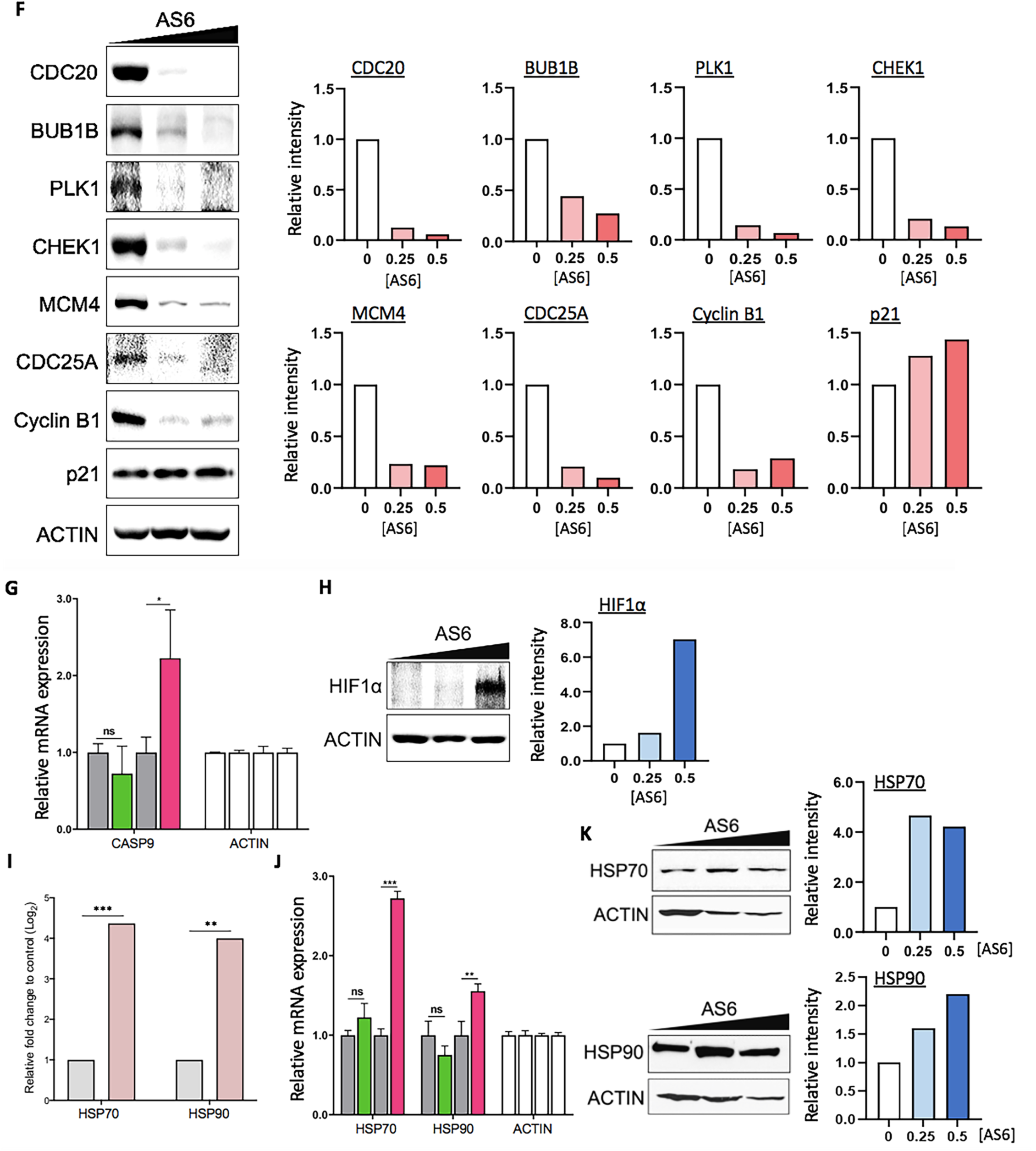
AS6 regulates DNA repair, cell cycle, and apoptosis in MCF7 cells. **(A)** Key DNA repair genes affected by AS6. RNA-seq data showing a markedly reduced expression of *ATM, p53*, and *BRCA1* in AS6-treated MCF7 cells. Light grey, untreated control; light pink, 0.5 μM AS6-treated cells. *n* = 3; * * P < 0.005; * * * P < 0.0005. **(B)** Real-time PCR data showing the expression of *ATM, p53*, and *BRCA1* in AS6-treated HUMEC and MCF7 cells. *ACTIN* was used as a reference gene for a normalization. From left to right bars, untreated control HUMEC, 0.5 μM AS6-treated HUMEC, untreated control MCF7 cells, 0.5 μM AS6-treated MCF7 cells. *n* = 3; error bars show s.d.; * * P < 0.005; * * * P < 0.0001. **(C)** Immunoblotting results showing a markedly reduced protein expression of ATM, p53, and BRCA1 in AS6-treated MCF7 cells. ACTIN was used as a loading control. Left, immunoblots; right, quantitative presentations of the signals, normalized with the control. AS6 concentrations in μM. Signal intensity of immunoblotting results throughout this manuscript was measured by Image J. Full-length blots were presented in Supplementary Fig. 4A. **(D)** Key cell cycle regulators affected by AS6. RNA-seq data showing a significantly reduced expression of *CDC20, BUB1B, PLK1, CHEK1, CDK1, MCM4*, and *CDC25A* in AS6-treated MCF7 cells but not in HUMEC. White, untreated control HUMEC; purple, 0.5 μM AS6-treated HUMEC; Light grey, untreated control MCF7 cells; light pink, 0.5 μM AS6-treated MCF7 cells. *n* = 3; * P < 0.01; * * P < 0.001; * * * P < 0.0001; * * * * P < 0.00001. **(E)** Real-time PCR results confirming the reduced mRNA expression of *CDC20, BUB1B, CHEK1, CDK1*, and *MCM4* in AS6-treated MCF7 cells but not in HUMEC. *ACTIN* was used as a reference gene for a normalization. From left to right bars, untreated control HUMEC, 0.5 μM AS6-treated HUMEC, untreated control MCF7 cells, 0.5 μM AS6-treated MCF7 cells. *n* = 3; error bars show s.d.; * P < 0.05; * * P < 0.005; * * * P < 0.0001. **(F)** Immunoblotting results demonstrating a dramatically disrupted protein expression of critical cell cycle regulators CDC20, BUB1B, PLK1, CHEK1, MCM4, CDC25A, cyclin B1, and p21 in AS6-treated MCF7 cells. ACTIN was used as a loading control. Left, immunoblots; right, quantitative presentations of the immunoblotting signals, normalized with the loading control. AS6 concentrations in μM. Full-length blots were presented in Supplementary Fig. 4B. **(G)** Key apoptotic factors affected by AS6. Real-time PCR results of *CASP9* showing its increased mRNA expression in MCF7 cells but not in HUMEC. *ACTIN* was used as a reference gene for a normalization. From left to right bars, untreated control HUMEC, 0.5 μM AS6-treated HUMEC, untreated control MCF7 cells, 0.5 μM AS6-treated MCF7 cells. *n* = 3; error bars show s.d.; * P < 0.05. **(H)** Immunoblotting showing a marked increase of HIF1*α* protein expression (left) and the quantification of the signal intensity (right) in AS6-treated MCF7 cells. ACTIN was used as a loading control. Full-length blots were presented in Supplementary Fig. 4C. **(I)** RNA-seq data of *HSP70* and *HSP90* mRNA expression in AS6-treated MCF7 cells. Light grey, untreated control; light pink, 0.5 μM AS6-treated cells. *n* = 3; * * P < 0.001; * * * P < 0.0001. **(J)** Real-time PCR results validating the increased *HSP70* and *HSP90* mRNA expression in AS6-treated MCF7 cells but not in HUMEC. From left to right bars, untreated control HUMEC, 0.5 μM AS6-treated HUMEC, untreated control MCF7 cells, 0.5 μM AS6-treated MCF7 cells. *n* = 3; error bars show s.d.; * * P < 0.005; * * * P < 0.0001. **(K)** Immunoblotting showing increased HSP70 and HSP90 protein expression (left) and the quantification of the signal intensity (right) in AS6-treated MCF7 cells. ACTIN was used as a loading control. Full-length blots were presented in Supplementary Fig. 4D.

Next, we examined and validated the genes known to regulate cell cycle progression that were differentially expressed between HUMEC and MCF7 cells in RNA-seq analysis (**Fig. 4D**). Of such genes, *CDC20*, BUB1 mitotic checkpoint serine/threonine kinase B (*BUB1B*), polo-like kinase 1 (*PLK1*), *CDK1*, checkpoint kinase 1 (*CHEK1*), mini-chromosome maintenance protein 4 (*MCM4*), and *CDC25A* were selected for validation through the real-time PCR and immunoblotting analysis (**Fig. 4E, F**; **Supplementary Fig. 4B**). CDC20 is an important regulator of cell division that activates anaphase promoting complex (APC/C) and also interacts with BUB1B [52, 53]. BUB1B encodes a kinase that functions in mitosis by inhibiting APC/C and ensuring kinetochore localization in CENPE and thus is required for a proper progression of mitosis[54]. PLK1 functions throughout the M phase of the cell cycle, phosphorylating many cell-cycle regulators, including BUB1B and cyclin B1 [55, 56]. PLK1 is important for the initiation of anaphase, the removal of cohesins, and spindle assembly [57]. CDK1 is a central cyclin-dependent kinase required for progression through the G_2_ and M phases [58]. CHEK1 is an essential protein that senses DNA damage to activate DNA repair machinery and cell cycle checkpoints throughout the cell cycle [59]. CDC25A is a phosphatase that removes the inhibitory phosphorylation from CDKs and controls the entry into the S and M phases [60]. MCM4 is a DNA helicase, an essential factor initiating genome replication in the S phase [61]. mRNA levels of these genes were compared in control and AS6-treated HUMEC and MCF7 cells. The results showed that mRNA expression of *CDC20, CDK1, MCM4, CHEK1*, and *BUB1B* was markedly reduced in MCF7 cells (**Fig. 4E**), which suggests a mechanism for the marked inhibition of cell cycle progression. By contrast, these mRNA levels were upregulated or less affected in HUMEC (**Fig. 4E**). These findings are consistent with the results of the cytotoxicity data (**Fig. 1**), which showed a deteriorated cell growth caused by AS6 in MCF7 cells. Immunoblotting results clearly showed that BUB1B, PLK1,CDC20, cyclin B1, CHEK1, and MCM4 were downregulated in AS6-treated MCF7 cells (**Fig. 4F**; **Supplementary Fig. 4B**). The mRNA expression of *CDC25A* was modestly increased in both HUMEC and MCF7 cells. However, its protein level was decreased in MCF7 cells (**Fig. 4E, F**; **Supplementary Fig. 4B**). Consistent with the RT-PCR results (**Fig. 1D, 4F**), protein levels of p21 were moderately increased in these cells. These data demonstrate that AS6 effectively deregulates the cell cycle in MCF7 cells. It potently interferes with the gene expression of central cell cycle regulators, important for the progression through the S and M phases, inhibiting the cell growth and proliferation in MCF7 cells.

A few genes that were differentially expressed in MCF7 cells are involved in the apoptotic pathway. These genes were *p53, p21, BUB1B, CASP9*, and hypoxia-inducible factor 1-alpha (*HIF1α*)[62-64]. Some of these genes, including *BUB1B, p53*, and *p21*, are also regulators of the cell cycle (**Fig. 4A– F**). The apoptotic pathway was shown to be stimulated in AS6-treated MCF7 cells but to be downregulated in HUMEC (**Fig. 2F, 3A**). The differential expression of these genes was validated through real-time PCR and Western blotting. mRNA expression of *CASP9* was markedly increased in MCF7 cells but not in HUMEC upon AS6 treatment (**Fig. 4G**). However, as discussed above, the expression of *p53* was decreased in the AS6-treated MCF7 cells (**Fig. 4A–C**). It appears that the decrease in *p53* could be anti-apoptotic [65, 66]. However, some studies indicate that a reduction or deficiency in p53 promotes genomic instability by compromising DNA double strand break repair and thus apoptosis [67], which suggests that the deregulation of p53, rather than an increase or decrease in expression, results in cell cycle arrest and apoptosis. As shown in **Figures 1D** and **4F**, in spite of the decrease in p53, the induction of p21 could be indicative of DNA damage and the onset of apoptosis in AS6-treated MCF7 cells [68]. The reduction in *BUB1B* mRNA and protein levels also supports an increase in chromosomal instability and apoptosis in MCF7 cells treated with AS6 (**Fig**. **4E, F**). In addition, immunoblotting showed a dramatic increase in HIF1*α* expression upon the treatment with AS6 (**Fig. 4H**; **Supplementary Fig. 4C**). HIF1*α* is activated by hypoxia and can trigger apoptosis for a prolonged hypoxic condition [63, 69]. HIF1*α* and heat-shock proteins have a close functional relationship. HIF1*α* induces the expression of HSP70 and HSP90 and these heat-shock proteins play an important role in stabilizing and degrading HIF1*α* [70]. In addition, the induction of HSP70 and HSP90 expression can indicate the cellular stress [71-73]. Therefore, we monitored mRNA and protein levels of *HSP70* and *HSP90* in AS6-treated MCF7 cells using RNA-seq, real-time PCR, and immunoblotting (**Fig. 4I–K**). Gene expression of *HSP70* and *HSP9*0 was increased in MCF7 cells upon AS6 treatment. By contrast, the transcription of these genes was not induced in HUMEC at the same concentrations of AS6 (**Fig. 4I, J**; **Supplementary Fig. 4D**). These data suggested that AS6 increases the cellular stresses in MCF7 cells but not in HUMEC at a certain range of concentrations.

## DISCUSSION

In this study, we used RNA-seq analysis to evaluate the effects of AS6 on genome-wide gene expression in human mammary epithelial cells for the first time to our knowledge. The effects of AS6 were compared in primary normal cells (HUMEC) and representative malignant cancer cells (MCF7cells). This work contributes to the field of AS6 chemotherapy by elucidating the medicinal effects of a chemical molecule through whole transcriptome analysis to better understand its molecular mechanisms and physiological impacts and by comparing the differential effects of this chemical in normal and cancerous mammary cells. Gene expression analyses provide essential information on the immediate and long-term effects of compounds on cells, which are the basic units of all life. We found that AS6 has distinctive gene expression profiles and cytotoxicity in these cancerous and normal breast epithelial cells at concentrations where AS6 differentially targets malignant cells, findings which are important to consider in developing a treatment for breast cancer based on AS6.

Our transcriptome analysis indicated that AS6 targets specific pathways in MCF7 cells concentrated on regulating the cell cycle, DNA repair, and apoptosis (**Fig. 3A–D**). The apoptotic pathways were upregulated whereas cell cycle and DNA repair mediators were downregulated in AS6-treated MCF7 cells (**Fig. 3A–D, 4A–F**). By contrast, AS6 suppressed the genes involved in apoptosis while activating the genes that regulate DNA repair and the cell cycle in HUMEC (**Fig. 2D–G**). In addition, the cytotoxicity analysis showed differential susceptibility to AS6 between these two cells, with an LC_50_ value for MCF7 cells about 10-fold lower than that for HUMEC (**Fig. 1**; **Table 1**). These results suggest more devastating effects of AS6 in cancer cells than in the normal mammary epithelial cells. In fact, we observed increased cell viability when HUMEC were treated with AS6 at concentrations up to 1 μM (**Fig. 1C**). The trend shown by our cytotoxicity data for HUMEC resembled the hormetic curve [74], a cellular biphasic response to a substance. It would be interesting, in future studies, to validate the differential sensitivity of normal and cancerous mammary cells by testing other primary and cancer cells and to understand what attributes to the difference in future.

The transcriptome study here showed that the DNA repair system, in particular, homologous recombination is markedly impaired in AS6-treated MCF7 cells. Moreover, the increased expression of HIF1*α* and heat shock proteins indicates that cells provoke stress responses under the influence of AS6 at given concentrations (**Fig. 4H–J**). We propose that the weakened homologous recombination is destructive to the cell cycle progression because of genomic instability [75, 76]. Improper DNA repair is likely to activate the cell cycle checkpoints, halts the cell cycle and to induce apoptosis in the AS6-treated MCF7 cells. In previous studies, AS6-mediated apoptosis was reported in other cell lines, including SW620 and MCF7 cells [4, 31, 77]. Those studies, which used targeted approaches, proposed an AS6-driven apoptosis through NF*κ*B inhibition and MAPK activation [4, 77]. The present study, which used an unbiased screening, suggests that the induction of apoptosis attributes to genomic instability, resulting from the severe reduction in key DNA repair enzymes, including BRCA1, ATM, p52, ATR, and RAD51 in AS6-treated MCF7 cells (**Fig. 3C, D, 4A–C**; **Table 3**). Mutations and deregulation of these enzymes have been reported in cancers [43, 47, 49-51, 76]. Therefore, the mechanism by which AS6 regulates the expression of these enzymes would be an important question to answer for future work.

The differential sensitivity to AS6 between HUMEC and MCF7 cells is note-worthy. At 0.5 μM AS6, HUMEC did not show malfunctions in DNA repair or cell cycle arrest while MCF7 cells did. AS6 is commonly poisonous to both cells at concentrations over 2 μM (**Fig. 1B, C, 5A**). We suggest that this critical difference in response at certain concentrations might be advantageous for treating breast cancers without affecting the normal cells and tissues. Arsenic trioxide is a successful anti-leukemia drug without serious side effects [24, 25]. This feature may be common to AS6 and arsenic trioxide. However, a transcriptome study that subjected the placenta to prenatal arsenic exposure found that somewhat different pathways were regulated [78], compared to those in the current study.

**Figure 5.**
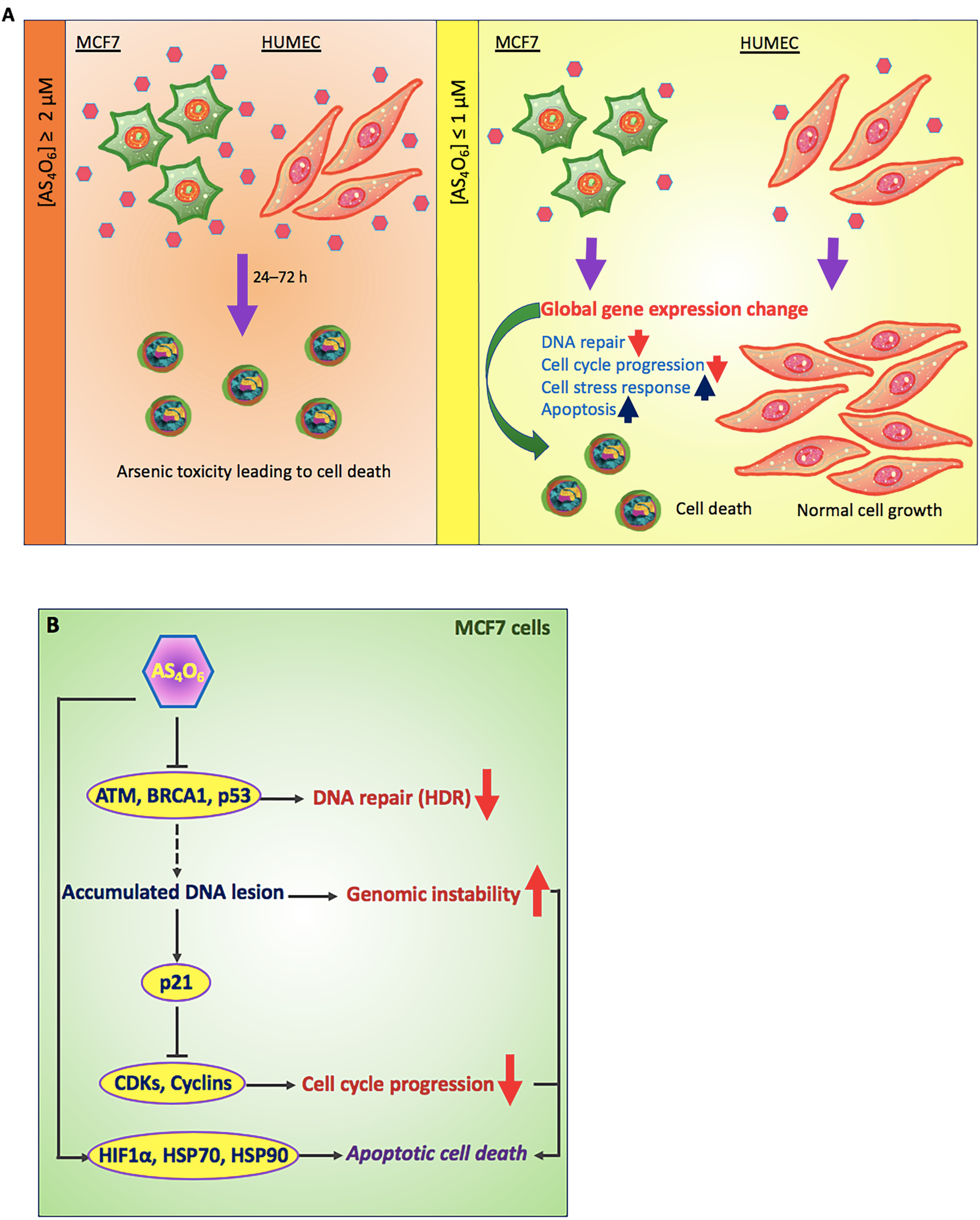
Proposed model of AS6 effects in normal mammary epithelial cells vs MCF7 cells. **(A)** In our study, it is shown that AS6 over 2 μM is toxic to both HUMEC and MCF7 cells to induce apoptotic cell death. By contrast, AS6 has cytotoxicity selectively to MCF7 cells, but not in HUMEC, at milder concentrations below 1 μM. Pinkish hexamers, AS6; rounded cells, apoptotic cells; upward arrow, upregulation; downward arrow; downregulation. **(B)** Proposed mode of action of AS6 (AS _4_O _6_). AS6 at milder concentrations might effectively target cancer cells to apoptotic cell death without much affecting normal cells. AS6 impairs the synthesis of key DNA repair enzymes for homologous recombination, which leads to genome instability. A potent CDKs/cyclins inhibitor p21 is activated to arrest the cell cycle while cell stress responses are provoked to facilitate the cell decision to apoptosis.

Comparison of this study with our current findings suggest a tissue-specific regulation by different arsenic compounds, which requires further study.

## CONCLUSIONS

Our novel findings suggest that AS6 at concentrations below 1 μM shows distinctive cytotoxicity and gene regulation profiles in a normal mammary epithelial and MCF7 cells: non-toxic and subtle changes in gene regulation in HUMEC but toxic and devastating alterations in gene regulation in MCF7 cells (**Fig. 5A**). The genomics data presented here suggest the mechanisms by which AS6 functions to suppress the proliferation of MCF7 cells. AS6 signaling obstructs the transcription of DNA repair enzymes whose function is crucial for homology-directed DNA double strand break repair. The genome becomes unstable, and this instability triggers the cell cycle arrest and apoptosis (**Fig. 5A, B**). In addition, AS6 provokes cell stress responses that promote apoptosis in MCF7 cells (**Fig. 5B**).

By contrast, the primary mammary epithelial cells show much greater resilience to AS6 at these same concentrations, in terms of the cellular stress level, cell growth, and the number of AS6-affected genes and -pathways (**Fig. 5A**). Therefore, we propose that AS6 below 1 μM induces cytotoxicity and derailed gene expression leading to cellular apoptosis in MCF7 cells, and potentially in malignant breast cancer cells, with milder impacts on surrounding normal breast cells.

## LIST OF ABBREVIATIONS

MCF7: Michigan Cancer Foundation 7
HUMEC: Human mammary epithelial cell
WHO: World Health Organization
ER: Estrogen receptor
APL: Acute promyelocytic leukemia
PML-RAR*α*: Promyelocytic leukemia protein-retinoic acid receptor *α*
HIV-1: Human immunodeficiency virus-1
DEGs: Differentially expressed genes
GO: Gene ontology
PPI: Protein-protein interaction
ATM: Ataxia-telangiectasia mutated
BRCA1: Breast cancer susceptibility 1
HDR: Homology-directed DNA double strand break repair
BUB1B: BUB1 mitotic checkpoint serine/threonine kinase B
PLK1: Polo-like kinase 1
CHEK1: Checkpoint kinase 1
MCM4: Mini-chromosome maintenance protein 4
APC/C: Anaphase promoting complex
HIF1*α*: Hypoxia-inducible factor 1*α*

## DECLARATIONS

### Ethic Approval and Consent to Participate

Not applicable

### Consent for Publication

Not applicable

### Availability of Supporting Data

All data are available in the manuscript or the supplementary material. The RNA-seq data has been deposited into NCBI Gene Expression Omnibus under accession number GSE157574.

### Authors’ Contributions

DK performed cell culture, cytotoxicity assays, quantitative real time PCR, mRNA quantification assays, statistical analysis, and Western blotting. NP and DC carried out cell culture, cytotoxicity assays, RNA preparation, and Western blotting. KK and HB performed bioinformatics analysis. SKC advised in the interpretation of the data. IB conceptualized and administrated the project, acquired funding, and provided with the materials. HB conceptualized and designed the experiments, prepared for the RNA samples, analyzed the data, and wrote the manuscript.

## Acknowledgement

We thank C. Li at Omega Bioservices (GA, USA) and Bunch lab members at Kyungpook National University (KNU) for their technical assistance and discussions. We appreciate X. Li at the University of Rochester Medical Center (NY, USA) and H. Cho at Korean Intellectual Property Office for the helpful discussion and mediating crucial collaboration for this study. H.B. thanks J. Christ and John and D. Y. Bunch for their loving encouragement throughout the course of this work.

